# Neuropeptides NLP-10 and FLP-18 stimulate lipid catabolism in chronically diet-restricted *Caenorhabditis elegans*

**DOI:** 10.64898/2026.09.17.752367

**Authors:** Jiaming Liu, Liyun Lin, Luming Yao, Yong Yu, Jintao Luo

## Abstract

Neuropeptides mediate inter-tissue communication in energy metabolism, yet most remain functionally uncharacterized. Using *C. elegans*, we uncover a neuropeptide signaling cascade that drives lipid catabolism under chronic dietary restriction. We show that impaired pharyngeal function leads to dyspepsia and nutrient insufficiency, which activates a signaling axis comprising two peptide-receptor pairs, NLP-10-NPR-35 and FLP-18-NPR-4. This cascade originates in ADL sensory neurons, relays through multiple interneurons, and enhances triglyceride hydrolysis in the intestine. Genetic disruption of any component of this axis restores fat accumulation in diet-restricted mutants, whereas overexpression of individual components in well-fed animals is insufficient to drive fat mobilization. Importantly, the same neuropeptide axis is engaged in starved wild-type animals, indicating a general response toward energy deficit rather than mutant-specific effects. These findings reveal a conserved neuroendocrine circuit that senses internal energy shortage and orchestrates systemic metabolic adaptation, providing a conceptual framework for how neuropeptide networks integrate nutritional status with energy balance in animals.

## INTRODUCTION

Energy metabolism is orchestrated by multi-tissue neuroendocrine networks in animals. Acting as messengers between the central nervous system and peripheral metabolic tissues, neuropeptides play key regulatory roles in this process, coupling nutrient availability to physiological responses that govern the storage or mobilization of neutral lipids. In mammals, hypothalamic and brainstem circuits integrate peripheral hormonal cues (e.g., insulin, leptin) with central neuropeptidergic signals (e.g., AgRP, α-MSH) to regulate adiposity and appetite^1,2^. Despite well-defined examples, the vast majority of neuropeptide systems remain functionally uncharacterized. In humans, although 462 neuropeptide-receptor pairs involving 225 neuropeptides have been predicted^3^, most have no characterized functions ^4^. This gap is further complicated by ligand-receptor redundancy and the often-dominant effects of hormones, which can obscure the subtle yet critical contributions of specific neuropeptide pathways to energy homeostasis^5–7^.

The nematode *Caenorhabditis elegans* (*C. elegans*)^8^ provides a powerful platform to dissect neuropeptide functions, given its evolutionarily conserved metabolic pathways^9^, genetically tractable neuropeptide connectomes^10,11^, and direct peptide-receptor communication via pseudocoelom^8^. A previous genetic screen revealed that mutants lacking functional Bone Morphogenetic Protein (BMP) signaling exhibit significant fat loss in intestine^12^, a phenotype confirmed in this study and by other publications^13,14^. Metabolic tracing demonstrated that the BMP-associated fat loss arises from accelerated lipid mobilization (e.g., mitochondrial β oxidation of fatty acids), rather than reduced fat synthesis^12^. The BMP signaling pathway is a branch of the TGF-β superfamily (**Figure 1—figure supplement 1A**)^15–18^; however, the mechanistic link between BMP deficiency and peripheral fat loss has remained elusive.

Notably, genetic restoration of BMP signaling in the pharynx fully rescued the intestinal fat accumulation^12^, exhibiting a cell-non-autonomous working pattern. Given that pharyngeal muscles of wildtype *C. elegans* provide food grinding forces to crush bacterial food^19^, and *dbl-1* mutants had more intestinal bacteria under normal feeding conditions^20,21^, we hypothesize that pharyngeal BMP signaling regulates food grinding to control intestinal fat storage via inter-tissue mechanisms. Long distance diffusion of neuropeptides allows inter-tissue communication^22–24^, serving as ideal candidates to mediate cell-non-autonomous responses.

Here, we uncovered that the BMP-associated fat loss is caused by pharyngeal impairments and subsequently insufficient food grinding, and the pharynx-to-intestine regulation relies on a hierarchy neuropeptide signaling axis. In this axis, the ADL sensory neurons release neuropeptide NLP-10 to activate its receptor NPR-35, which functions in interneurons to promote the secretion of a lipid-regulating neuropeptide, FLP-18. Abundant FLP-18 stimulates the intestinal receptor NPR-4 to promote fat loss, a process requiring adipose triglyceride lipase (ATGL-1) and other lipases. Importantly, this neuropeptide axis is not merely a mutant-specific phenomenon: In fasted wild-type animals, the same NLP-10/NPR-35 and FLP-18/NPR-4 modules are engaged to promote lipid catabolism, suggesting a general, conserved neural mechanism for fat mobilization in response to nutrient deprivation. Our findings establish neuropeptides as systemic neuroendocrine hubs that link nutrient processing to organism-wide metabolic adaptation, thereby synchronizing animal physiology with nutrient availability.

## RESULTS

### BMP mutants are dyspeptic and chronically diet-restricted

Consistent with previous findings^12^, suppression of DBL-1 or SMA-6 was sufficient to reduce the intestinal fat contents in *C. elegans* (**Figure 1A, Figure 1—figure supplement 1B**). By scoring colony-forming units (CFU) within the *C. elegans* gastrointestinal tract, and by measuring relative fluorescent intensity of the GFP-positive bacterial food (OP50-GFP) within *C. elegans’* gastrointestinal tract, we affirmed that the BMP mutants contained more viable bacteria (**Figure 1B, Figure 1—figure supplement 1C**). Observation of intestinal cross-sections by transmission electron microscopy (TEM) revealed frequent bacterial presence in the gastrointestinal lumen of the BMP mutants, but not in wildtype (**Figure 1C**, **Figure 1— figure supplement 1D**).

**Figure 1.**
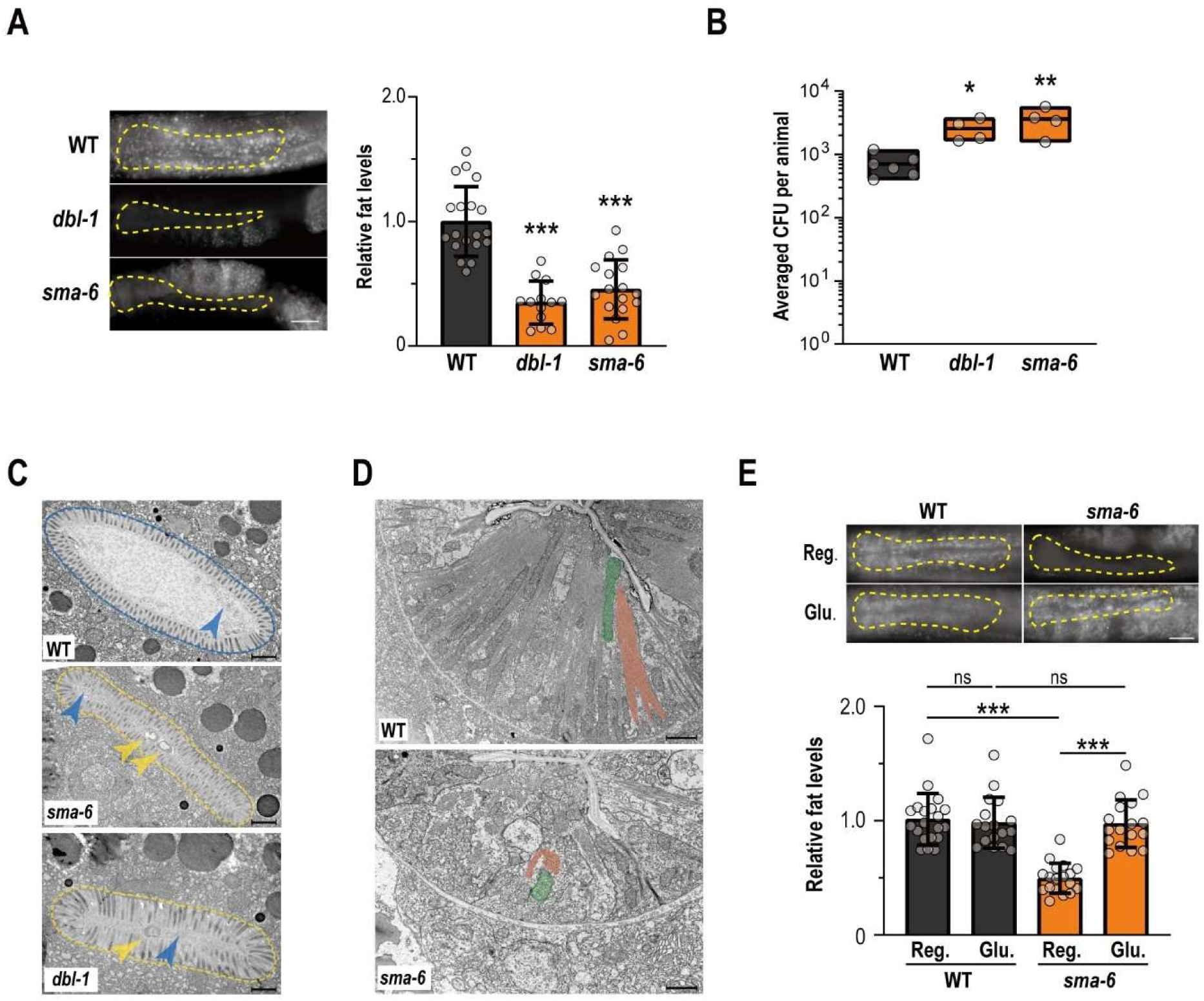
Pharyngeal dysfunction leads to dyspepsia in BMP mutants. (**A**) Relative fat levels and representative images of wildtype, *dbl-1(wk70),* and *sma-6(wk7).* (**B**) Averaged colony-forming units (CFU) in intestines of wildtype, *sma-6(wk7)* and *dbl-1(wk70)*. (**C**) Transmission electron microscopy (TEM) images of wildtype, *sma-6(wk7)* and *dbl-1(wk70)* at the animals’ intestine. The yellow arrows indicate bacteria, and the blue arrows indicate bacterial lysate. (**D**) TEM images of wildtype and *sma-6(wk7)* at the animals’ pharynx. The green shade indicates one mitochondrion, and the pink shade indicates a bundle of myofilaments. (**E**) Relative fat levels and representative images of wildtype and *sma-6(wk7)* feeding on the regular OP50 diet or the regular diet supplemented with glucose. In **A**, **B**, **E**, the Kruskal-Wallis tests with Dunn’s multiple comparison between all groups were applied, with * indicates *p* < 0.05, ** indicates *p* < 0.01, *** indicates *p* < 0.001, and ‘ns’ indicates not significant. In **A** and **E**, all representative images in the same panel share the same scale indicated by a 25 μm scale bar. In **C** and **D**, all scale bars represent 1 μm.

Pharyngeal muscles of wildtype *C. elegans* are fulfilled with myofilaments aligned at angles approximately perpendicular to the anterior-posterior body axis^25^. Indeed, cross-section TEM images of wildtype pharynx showed long and continuous myofilaments accompanied by healthy mitochondria; In contrast, myofilaments of *sma-6(wk7)* are short, gapped, and chaotically aligned, accompanied by pale mitochondria (**Figure 1D**). Above findings suggest that lacking BMP pathway has impaired the pharyngeal muscle, which could very likely reduce the pharyngeal food grinding force. Consistently, previous findings confirmed the requirement of pharyngeal BMPs in regulating intestinal fat storage^26^. Simultaneously, we detected bacterial lysate (similar to the lysate of ultrasound-treated *E. coli*^27^) in the gastrointestinal lumens of both wildtype and BMP mutants (**Figure 1C**). Although we did not distinguish whether bacterial death occurred prior to ingestion or resulted from pharyngeal grinding, we think that the BMP mutants could derive at least some nutrients (if not sufficient amounts) from their routine laboratory diet.

If nutrient deficiency was the reason behind BMP-associated fat loss, then we would expect the BMP mutants to benefit from absorbing edible nutrients for fat accumulation. We respectively supplemented wildtype or *sma-6(wk7)* with small molecule nutrients – either glucose, fructose or oleic acid (OA) – in addition to the routine bacterial diet. All three types of nutrients restored the intestinal fat levels in *sma-6(wk7)* at relatively low doses that would not disturb wildtype fat contents (**Figure 1E, Figure 1—figure supplement 1E, F**). Such nutrient-dependent fat restoration likely occurred via metabolic pathways downstream of or independent from the defective BMP signaling, demonstrating that the BMP mutants are viable and fertile animal models with consistent dyspepsia and chronical dietary restriction.

### Neuropeptide signaling promotes lipid catabolism in *sma-6* mutants

By analyzing the relative gene expression levels of *sma-2(rax5)* versus wildtype, and *sma-4(rax10)* versus wildtype^12^, we predicted differentially expressed genes (DEGs). Then we utilized Gene Ontology (GO) analysis on the intersection of genes induced in both *sma-2(rax5)* and *sma-4(rax10)*, and the neuropeptide signaling pathway was highlighted (**Figure 2—figure supplement 1A**). Given that the secretion of neuropeptides require dense core vesicles (DSVs), we disrupted *unc-31*, which encodes the conserved Ca^2+^-dependent activator protein for secretion (CAPS) to mediate DSV docking and priming in *C. elegans* (**Figure 2—figure supplement 1B**)^28–30^. Disruption of *unc-31* facilitated intestinal fat accumulation in BMP mutants (**Figure 2—figure supplement 1C**), confirming the involvement of neuropeptide secretion in regulating BMP-associated fat loss.

To address the specific neuropeptides in regulating fat mobilization, we verify the RNA-seq results by qRT-PCR on total mRNA samples from young adults of wildtype, *sma-6(wk7), dbl-1(wk70)* and *sma-2(rax5)*, with primers respectively targeting neuropeptide-encoding genes (**Figure 2—figure supplement 2D, supplementary Table 1**). Besides, we compared our qRT-PCR results (rectangles) to the 16h-fasted gene expression data in Harvald et al^31^ (ovals). Indeed, a majority of neuropeptide genes exhibited upregulation in both datasets (**Figure 2—figure supplement 1D**).

Next, we obtained either a loss-of-function mutant or a feeding-based RNAi strain for 41 of 73 neuropeptide genes. For each, we generated double knockdowns of a neuropeptide gene and a BMP gene. Then we compared the intestinal fat levels of the double knockdowns to their corresponding siblings with functional neuropeptides but dysfunctional BMP signaling (**Figure 2—figure supplement 1E, supplementary Table 2**). Among all genes required in the BMP-associated fat loss, despite that some neuropeptides were known to regulate lipid metabolism under other contexts (e.g., *flp-18*^24,32^ and *nlp-12*^33^), the evolutionarily conserved *nlp-10* attracted our attention, as it was consistently upregulated in different types of BMP mutants and starved animals, and also capable to suppress fat accumulation in the BMP mutants (**Figure 2A, Figure 2—figure supplement 1E, F**). Furthermore, we generated transgenic strains overexpressing *nlp-10* by an endogenous promoter or a pan-neural promoter in *sma-6(wk7); nlp-10(tm6232)*, and these transgenic animals failed to accumulate sufficient fat, phenocopying the low-fat status of *sma-6(wk7)* (**Figure 2B, C**).

**Figure 2.**
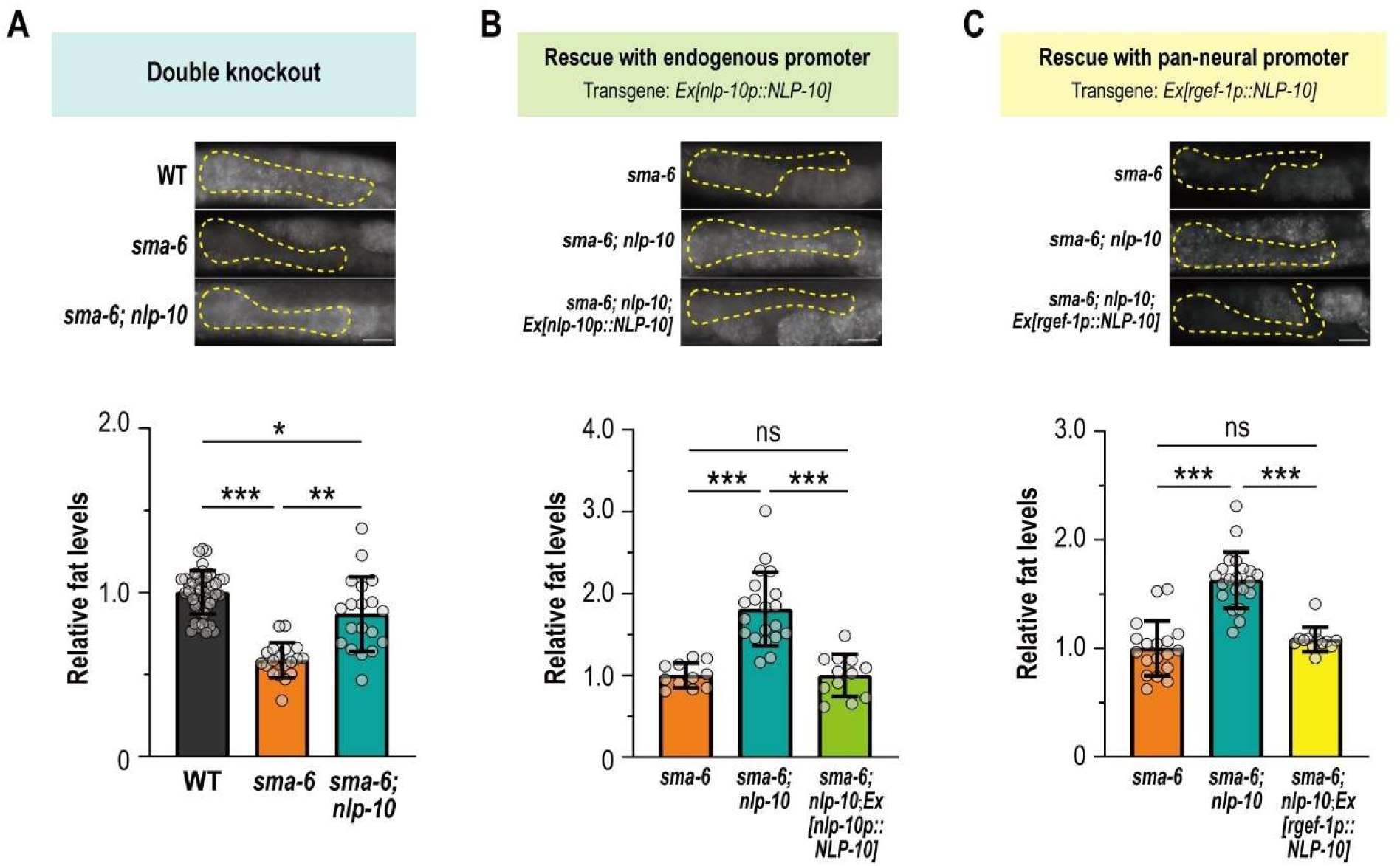
NLP-10 is required for the BMP-associated fat loss. (**A**) Relative fat levels and representative images of wildtype, *sma-6(wk7)*, and *sma-6(wk7); nlp-10(tm6232)*. (**B-C**) Relative fat levels and representative images of *sma-6(wk7)*, *sma-6(wk7); nlp-10(tm6232)*, and *sma-6(wk7); nlp-10(tm6232)* with transgenes expressing *nlp-10* by the endogenous promoter (**B**) or the pan-neural promoter (*rgef-1p*) (**C**). For **A-C**, the Kruskal-Wallis test and Dunn’s multiple comparison between all groups were applied, with * indicates *p* < 0.05, ** indicates *p* < 0.01, *** indicates *p* < 0.001, and ‘ns’ indicates not significant. For **A-C**, all representative images in the same panel share the same scale indicated by a 25 μm scale bar.

### BMP-associated fat loss is regulated by NLP-10 secreted from ADL neurons

Based on *C. elegans* gene expression databases including WormSeq^34^, CeNGEN^35^, CAWA^36^, and CaenoGen^37^, as well as the DiI-based imaging on the transcriptional reporter *nlp-10p::GFP*, we confirmed that *nlp-10* mainly expresses in neurons, including amphid neurons ADL (**Figure 3A**). In *sma-6(wk7); nlp-10(tm6232)* with ADL-specific expression of *nlp-10*, the fat levels were as low as that of *sma-6(wk7)* (**Figure 3B**). Consistently, genetic ablation of ADL neurons in *sma-6(wk7)*, achieved by two independent ADL-specific TeTx (light chain of tetrodotoxin) lines^38^, enabled fat restoration in *sma-6(wk7)* (**Figure 3C**), resembling the *nlp-10* suppression in BMP mutants.

**Figure 3.**
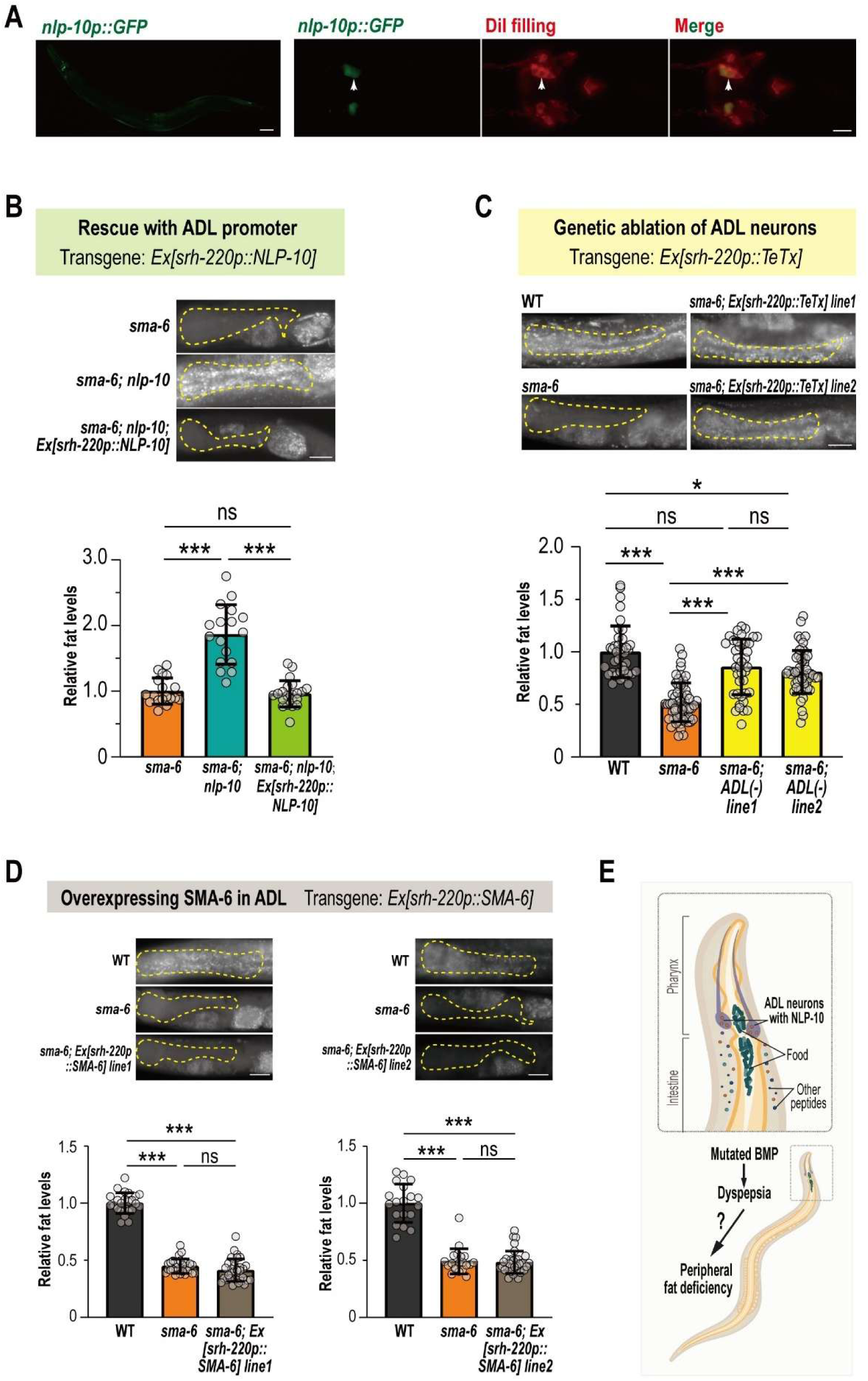
NLP-10 but not SMA-6 is required for the BMP-associated fat loss in the ADL neurons. (**A**) The very left image indicates the fluorescence of a transcriptional reporter *nlp-10p::GFP* in a whole worm (scale bar = 25 μm), the other images indicate fluorescence of the same *nlp-10p::GFP* reporter (green) labeled by DiI staining (red) with the white arrows indicating an ADL neuron (scale bar = 10 μm). (**B**) Relative fat levels and representative images of *sma-6(wk7)*, *sma-6(wk7); nlp-10(tm6232)*, and *sma-6(wk7); nlp-10(tm6232)* with transgenes expressing *nlp-10* by the ADL-specific promoter (*srh-220p*). (**C**) Relative fat levels and representative images of wildtype, *sma-6(wk7)* and ADL-ablated *sma-6(wk7)* lines achieved by transgenic *srh-220p::TeTx*. (**D**) Relative fat levels and representative images of wildtype, *sma-6(wk7)*, and *sma-6(wk7)* with transgenes expressing *sma-6* by the ADL-specific promoter (*srh-220p*). (**E**) A model demonstrating a hypothesis that BMP mutation leads to pharyngeal impairment and thus dyspepsia, which ultimately causes peripheral fat deficiency, and the NLP-10 neuropeptides released from ADL neurons are required to mediate the fat loss.

The pharyngeal rescue of SMA-3 by the *myo-2p::SMA-3* transgene has been shown to restore the low-fat phenotype of *sma-3(wk30)* ^26^, and ADL neurons are known to express *sma-6* in addition to its *nlp-10* expression^34,35^. So, we wondered if *sma-6* might have worked in the ADL neurons to transcriptionally regulate *nlp-10*, and we specifically restored the expression of SMA-6 in the ADL neurons of *sma-6(wk7)*. Yet the fat levels remained low in these transgenic worms (**Figure 3D**), suggesting that *sma-6* does not work in the ADL neurons to regulate fat contents. Together, we hypothesize a model that the BMP deficiency has impaired pharyngeal food grinding to generate chronical dietary-restriction at individual level, and such diet-restricted status would trigger the ADL neurons to release NLP-10 to remotely facilitate lipid catabolism in intestine (**Figure 3E**). In BMP mutants, given that suppressing other peptides was also sufficient to alleviate fat loss (**Figure 2—figure supplement 1E**), NLP-10 is unlikely to be the exclusive peptide with lipid regulatory roles. Yet it serves as a good entry point for us to further investigate the downstream mechanisms, as it is robustly upregulated in all examined BMP mutants, and also has a known receptor, NPR-35^11,39^. Based on these findings, we sought to focus on how NLP-10 mediates lipid catabolism.

### NPR-35 functions in interneurons to promote BMP-associated fat loss

To our knowledge, *npr-35* encodes the sole known receptor of NLP-10^11,39–42^. Fat levels of *sma-6(wk7); npr-35(ok3258)* were greater than that of *sma-6(wk7)*, resembling *sma-6(wk7); nlp-10(tm6232)*; However, the triple mutant *sma-6(wk7); nlp-10(tm6232); npr-35(ok3258)* accumulated even more fat than that of *sma-6(wk7); nlp-10(tm6232)* or *sma-6(wk7); npr-35(ok3258)* (**Figure 4A**), indicating involvement of unknown ligands for *npr-35* or unknown receptors for *nlp-10*, or both.

**Figure 4.**
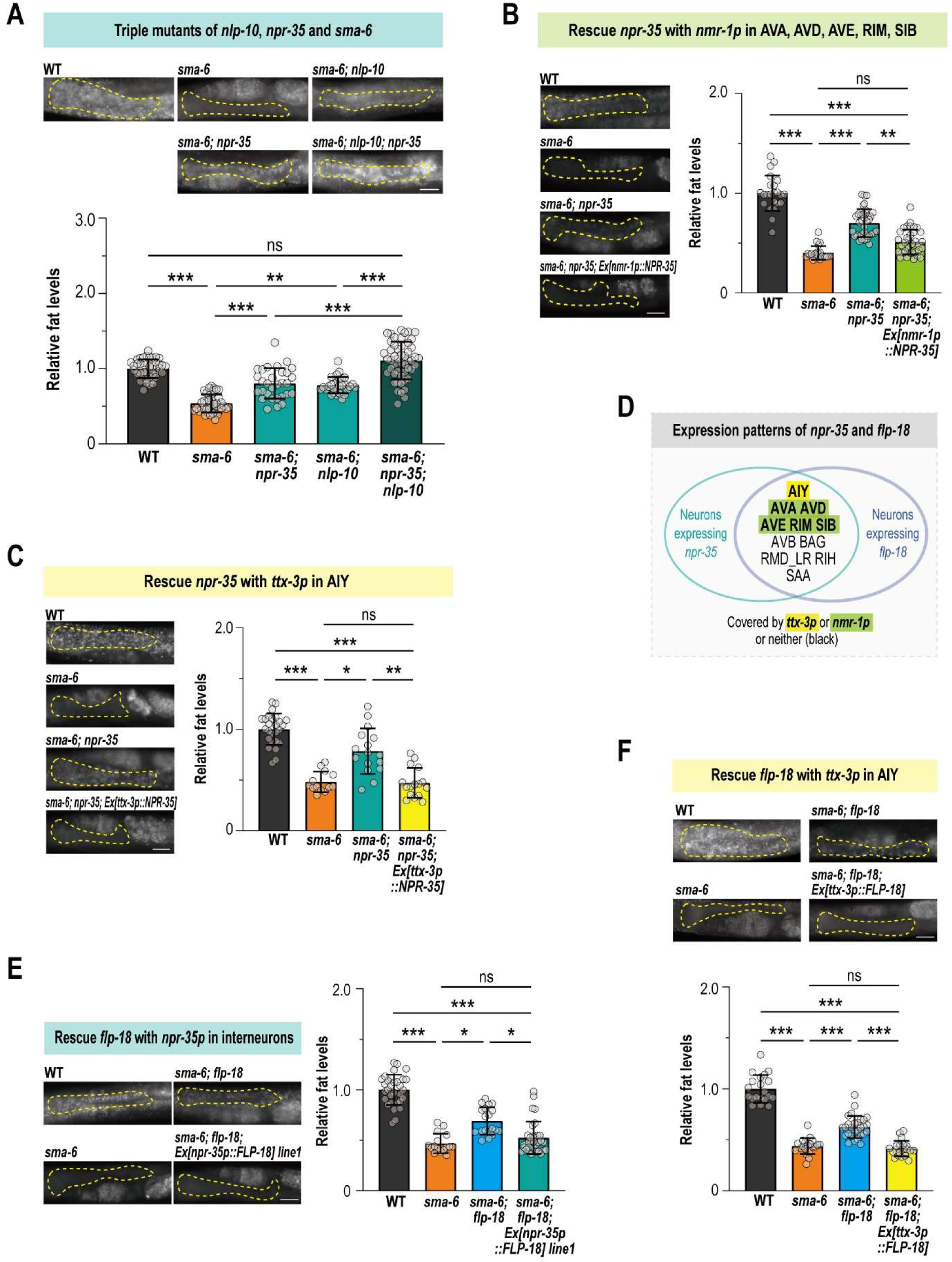
Neuropeptide receptor NPR-35 and neuropeptide FLP-18 mediates fat loss in BMP mutants. (**A**) Relative fat levels and representative images of wildtype, *npr-35(ok3258)*, *sma-6(wk7)*, *sma-6(wk7); npr-35(ok3258)*, *sma-6(wk7); nlp-10(tm6232)* and *sma-6(wk7); npr-35(ok3258); nlp-10(tm6232)*. (**B-C**) Relative fat levels and representative images of wildtype, *sma-6(wk7)*, *sma-6(wk7); npr-35(ok3258)*, and *sma-6(wk7); npr-35(ok3258)* with transgenes expressing *npr-35* by the interneuron promoter (*nmr-1p*) (**B**) or the AIY promoter (*ttx-3p*) (**C**). (**D**) A Venn’s diagram demonstrates the expressive intersections of *npr-35* and *flp-18*. (**E-F**) Relative fat levels and representative images of wildtype, *sma-6(wk7)*, *sma-6(wk7); flp-18(tm2179)* and *sma-6(wk7); flp-18(tm2179)* with *flp-18* expression in the *npr-35-*expressing neurons (**E**) or with *flp-18* expression in AIY (**F**). For **A**, **B**, **C**, **E** and **F**, the Kruskal-Wallis test and Dunn’s multiple comparison between all groups were applied, with * indicates *p* < 0.05, ** indicates *p* < 0.01, *** indicates *p* < 0.001, and ‘ns’ indicates not significant. For **A**, **B**, **C**, **E** and **F**, all representative images in the same panel share the same scale indicated by a 25 μm scale bar.

The *npr-35*-expressing tissues mainly covers interneurons^34,35,43^, including AIY, AVA, AVD, AVE, etc. In *sma-6(wk7); npr-35(ok3258)*, we performed AVA, AVD, AVE, RIM, SIB-specific rescue of *npr-35* with *nmr-1* promoter (**Figure 4B**), and AIY-specific rescue of *npr-35* with *ttx-3* promoter (**Figure 4C**), and fat levels of *sma-6(wk7); npr-35(ok3258)* were suppressed by either transgene, supporting ectopic regulation of NPR-35 on BMP-associated fat deficiency. Interestingly, removal of *npr-35* failed to offset the fat deficiency in *sma-6(wk7); Ex[nlp-10p::NLP-10]* (**Figure 4—figure supplement 1A**), further suggesting that one or more unrevealed NLP-10 receptors could be required in regulating BMP-associated fat loss. Meanwhile, the neuronal but not gastrointestinal function of NPR-35 indicates that downstream mechanisms, which should signal from interneurons to intestine, are required to regulate BMP-associated fat loss.

### *npr-35*-expressing neurons release FLP-18 to mediate fat mobilization in BMP mutants

Interestingly, a large proportion of *npr-35-*expressing neurons also express *flp-18* (**Figure 4D**), a peptide that is required for the BMP-associated fat loss in our initial screens (**Figure 2—figure supplement 1D, E**). This expressive intersection has included AIY, AVA, AVD and other aforementioned interneurons^34,35^. So, we went on to investigate links between NLP-10, NPR-35 and FLP-18.

To verify if the *npr-35-*expressing neurons would alter fat contents via FLP-18, we compared the fat levels of *sma-6(wk7); flp-18(tm2179)* strains with or without the *npr-35p::FLP-18* transgene, and it turns out that the fat contents of *sma-6(wk7); flp-18(tm2179)*; *Ex[npr-35p::FLP-18]* was as low as that of *sma-6(wk7)* (**Figure 4E, Figure 4—figure supplement 1B**). Besides, expression of *flp-18* in the AIY neurons achieved by the *ttx-3p::FLP-18* transgene in *sma-6(wk7); flp-18(tm2179)* could restore fat levels to be *sma-6(wk7)*-like (**Figure 4F**). Together, we conclude that *flp-18* was required in *npr-35*-expressing interneurons for the *sma-6(wk7)* animals to mobilize fat.

### Downstream of FLP-18, intestinal NPR-4 mediates BMP-associated fat loss

As a known receptor of FLP-18, NPR-4 regulates intestinal fat storage in other contexts^24,32^. A transcriptional reporter of *npr-4p::GFP* confirmed its neuronal and intestinal expression (Figure 5A), and knockout of *npr-4* in BMP mutants led to restored fat accumulation (**Figure 5B**). Genetic rescue of *npr-4* by its endogenous promoter (**Figure 5C**) or the intestinal-specific promoter *ges-1p* (**Figure 5D, Figure 5— figure supplement 1A**) could suppress fat accumulation in *sma-6(wk7); npr-4(tm1782)*, supporting an *in-situ* lipid regulatory role of NPR-4 in BMP mutants.

**Figure 5.**
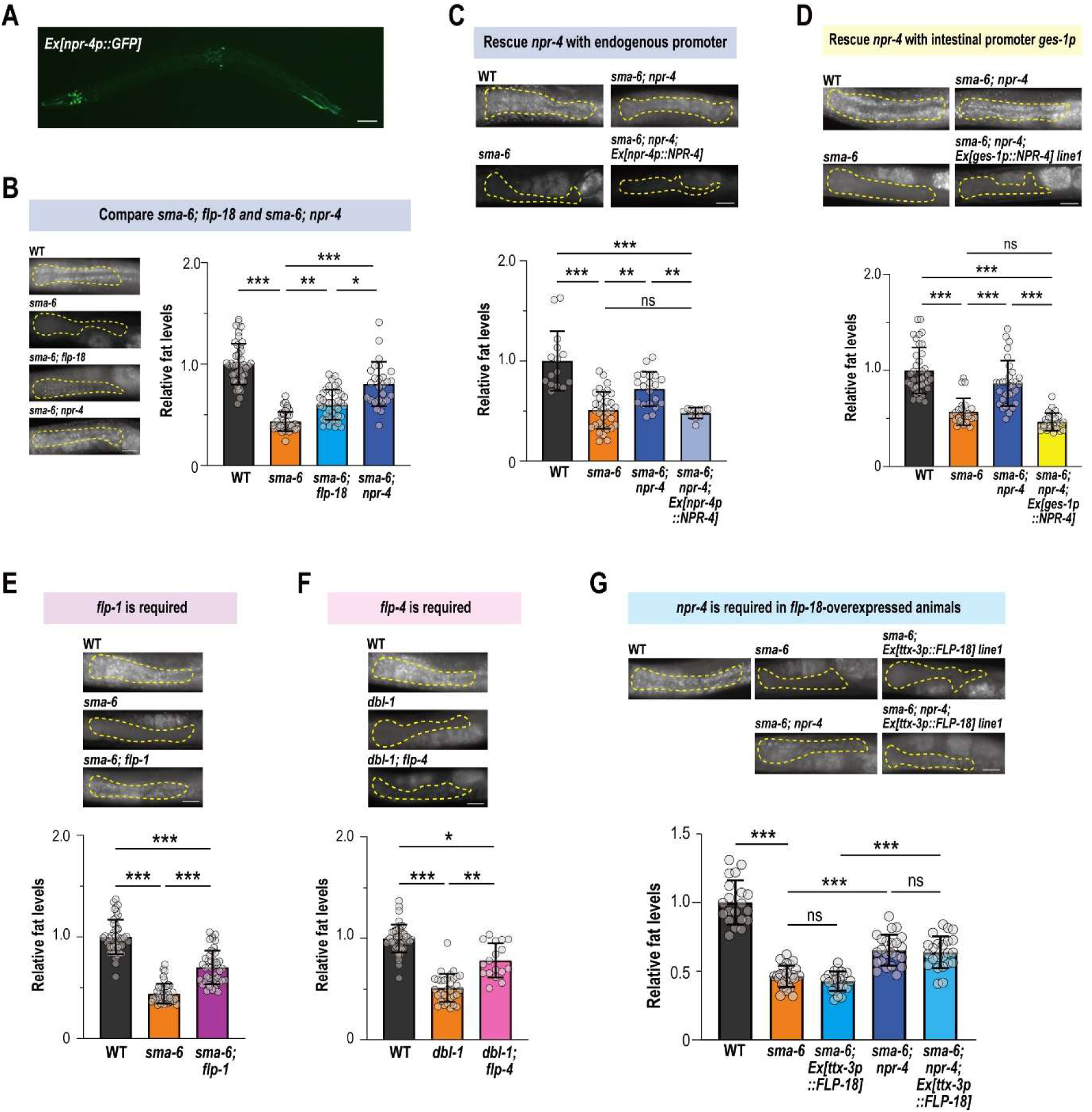
Intestinal NPR-4 is required for the BMP-associated fat deficiency. (**A**) Fluorescence of a transcriptional reporter *npr-4p::GFP* (scale bar = 50 μm). (**B**) Relative fat levels and representative images of wildtype, *sma-6(wk7)*, *sma-6(wk7); flp-18(tm2179)* and *sma-6(wk7); npr-4(tm1782)*. (**C-D**) Relative fat levels and representative images of wildtype, *sma-6(wk7)*, *sma-6(wk7); npr-4(tm1782)*, and *sma-6(wk7); npr-4(tm1782)* with *npr-4* expressed by the endogenous promoter (**C**) or the intestinal promoter *ges-1p* (**D**). (**E**) Relative fat levels and representative images of wildtype, *sma-6(wk7)*, and *sma-6(wk7); flp-1(sy1599)*. (**F**) Relative fat levels and representative images of wildtype, *dbl-1(wk70)*, and *dbl-1(wk70); flp-4(sy1606)*. (**G**) Relative fat levels and representative images of wildtype, *sma-6(wk7)*, *sma-6(wk7); npr-4(tm1782*), and *sma-6(wk7)* or *sma-6(wk7); npr-4(tm1782)* with *nlp-10* overexpressed by the endogenous promoter. For **B-G**, the Kruskal-Wallis test and Dunn’s multiple comparison between all groups were applied, with * indicates *p* < 0.05, ** indicates *p* < 0.01, *** indicates *p* < 0.001, and ‘ns’ indicates not significant. For **B-G**, all representative images in the same panel share the same scale indicated by a 25 μm scale bar.

Notably, *sma-6(wk7); npr-4(tm1782)* contained more fat than that of *sma-6(wk7); flp-18(tm2179)* (**Figure 5B**), indicating ligand redundancy. We tested other known ligands of NPR-4^39,40^, and we found that FLP-4 and FLP-1 were also required to mediate BMP-associated fat loss (**Figure 5E-F**), whereas FLP-14 and FLP-21 were dispensable (**Figure 5—figure supplement 1B, C**). Unlike FLP-18, which is co-expressed with NPR-35, FLP-1 has very limited expressive overlap with this receptor^34,35^, and our data demonstrate that FLP-1 does not seem to be secreted from the *npr-35-expressing* neurons (**Figure 5—figure supplement 1D**).

To test if *npr-4* deletion was sufficient to block the BMP-associated fat loss mediated by FLP-18, we directly compared the fat levels of *sma-6(wk7); npr-4(tm1782); Ex[ttx-3p::FLP-18]* with *sma-6(wk7); npr-4(tm1782)* and *sma-6(wk7); Ex[ttx-3p::FLP-18]*. Regardless of FLP-18 overexpression, *sma-6(wk7); npr-4(tm1782)* contained equivalently abundant amount of fat (**Figure 5G**), suggesting that *npr-4* works downstream of *flp-18*. On the other hand, overexpression of the upstream neuropeptide NLP-10 cannot alter the fat levels of *sma-6(wk7); npr-4(tm1782)*, either (**Figure 5—figure supplement 1E**). These results imply a regulatory axis consisted of NLP-10-to-NPR-35, FLP-18-to-NPR-4 in the upstream-to-downstream order, with FLP-1 and FLP-4 being additively required in parallel to FLP-18.

### Fat breakdown in BMP mutants requires lipolysis

By lipidomic analysis, in *sma-6(wk7)* we found that a majority of triglycerides (TG) were reduced, and almost all diglycerides (DG) were increased (**Figure 6—figure supplement 1, supplementary table 3**), and the TG-to-DG ratio was significantly lower in *sma-6(wk7)*, demonstrating enhanced triacylglycerol hydrolysis in these mutants. Suppressing the lipases’ activity in *sma-6(wk7)*, by RNAi of *atgl-1*, *lips-3, 4, 5* or *17*^44^, facilitated fat restoration (**Figure 6B-C**), supporting that the active fat breakdown in BMP mutants is driven by enhanced lipolysis.

**Figure 6.**
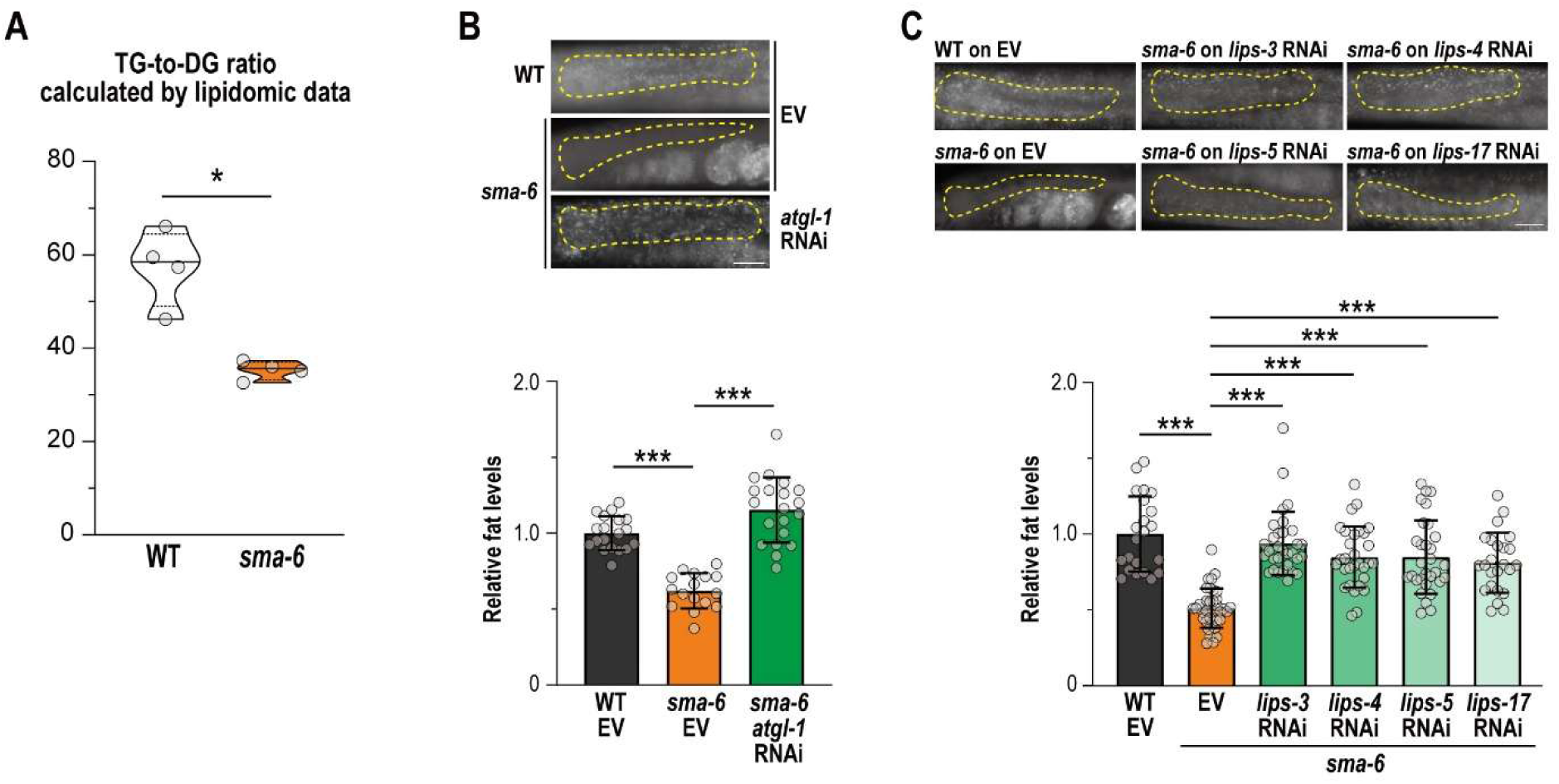
Lipases are required for the BMP-associated fat loss. (**A**) The TG-to-DG ratio calculated by lipidomic data in wildtype and *sma-6(wk7)*, related to **supplementary table 3**. (**B-C**) Relative fat levels and representative images of wildtype and *sma-6(wk7)* feeding on the *atgl-1* RNAi bacteria (**B**) or the *lips-3*,*4*,*5, 17* RNAi bacteria (**C**), controlled by the bacteria containing an empty vector L4440 (EV). The Mann-Whitney’s test was applied to **A**, and the Kruskal-Wallis test and Dunn’s multiple comparison between all groups were applied to **B** and **C**. For all statistical assays, * indicates *p* < 0.05, *** indicates *p* < 0.001, and ‘ns’ indicates not significant. In **B** and **C**, all representative images in the same panel share the same scale indicated by a 25 μm scale bar.

### NLP-10 and FLP-18 are starvation-induced neuropeptides

As hypothesized in the **figure 3E** model, if the peripheral fat loss was triggered by dietary restriction *per se*, then we would expect the same neuropeptides being required in other types of diet-restricted worms, but not merely dyspepsia-induced dietary restriction. To investigate whether this neuropeptide axis was generally required in diet-restricted animals or specifically required in BMP-deficient animals, we starved wildtype and single mutants of *nlp-10*, *flp-18*, *npr-35* or *npr-4* before measuring fat levels. 24 hours of starvation halved intestinal lipid content in wildtype, while each single mutant maintained higher fat levels under the same starved condition (**Figure 7A**), phenocopying well-fed double mutants such as *sma-6(wk7); nlp-10(tm6232)* and so on. By qRT-PCR, we demonstrated that the expression of *nlp-10*, *flp-18*, *npr-35* and *npr-4* were induced in starved wildtype and starved BMP mutants (**Figure 7B**), suggesting that the induction of *nlp-10* and *flp-18* in the BMP mutants are more likely to be triggered by a general energy-insufficient status, rather than a BMP-specific regulation. In *eat-4(ky5)* and *eat-2(ad1116)*, which are BMP-independent models with restricted energy intake^45,46^, we found that *flp-18* and *nlp-10*, as well as several other previously screened neuropeptides that may not mediate BMP-associated fat loss (encoded by *nlp-15*, *21*, *51* and *flp-14*, *27*) (**Figure 2—figure supplement 1D, E**), were upregulated (**Figure 7C**). This further supports NLP-10 and FLP-18 as general starvation-induced neuropeptides, and also verifies the findings of Harvald et al^31^. Collectively, although dozens of neuropeptides are transcriptionally induced by energy deficiency, only a subset of them are genetically required to stimulate fat mobilization.

**Figure 7.**
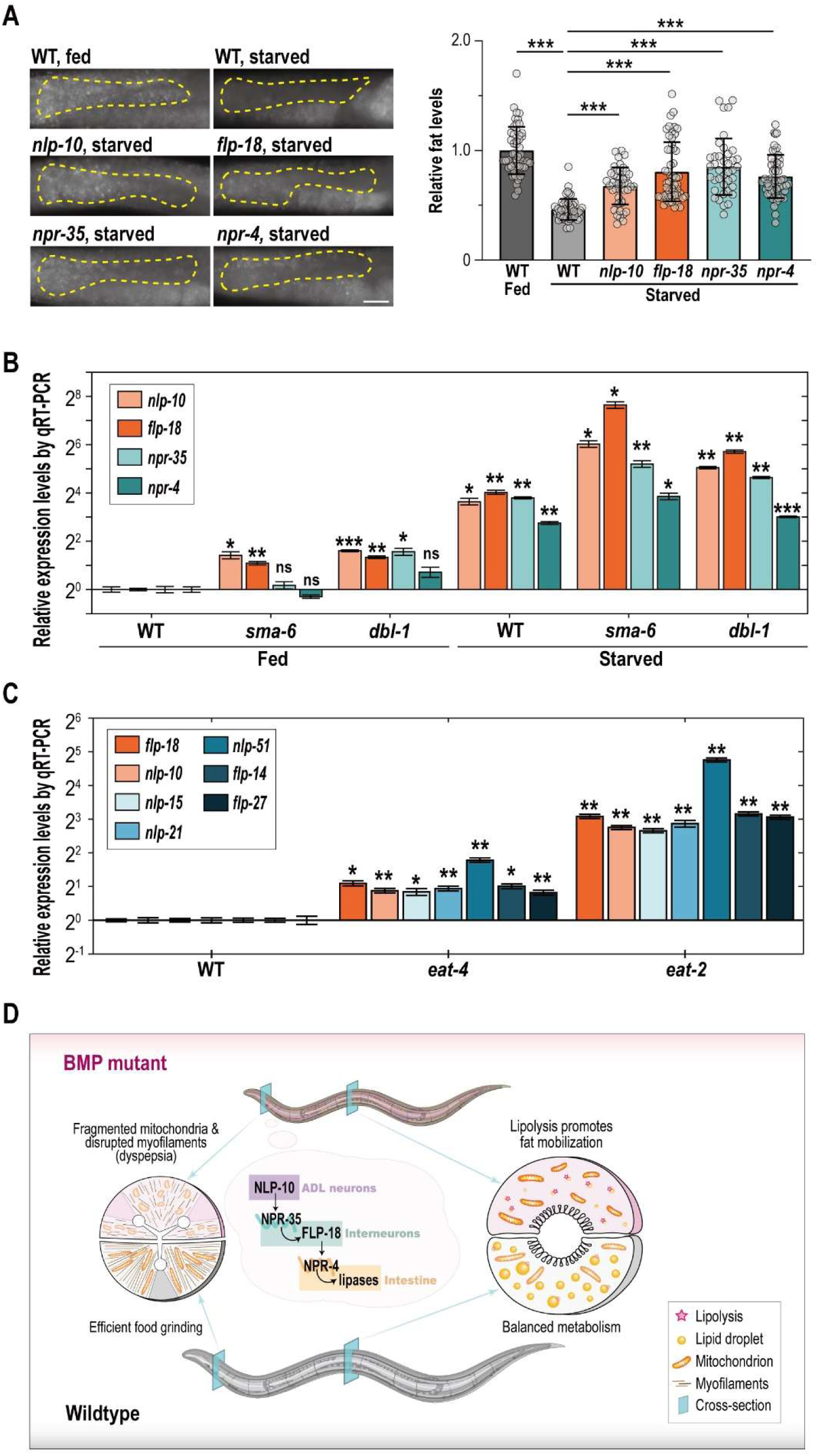
The neuropeptide axis responds to general energy deficiency. (**A**) Relative fat levels and representative images of wildtype and starved wildtype, *nlp-10(tm6232)*, *flp-18(tm2179)*, *npr-35(ok3258)* or *npr-4(tm1782)*. (**B**) Transcriptional levels of *nlp-10*, *flp-18*, *npr-35* and *npr-4* in wildtype or *sma-6(wk7)* or *dbl-1(wk70)* at well-fed or starved status, determined by qRT-PCR (the 2-ΔΔCt method). (**C**) Transcriptional levels of *flp-18*, *nlp-10*, *nlp-15*, *nlp-21*, *nlp-51*, *flp-14* and *flp-27* in wildtype or *eat-4(ky5)* or *eat-2(ad1116)* at well-fed status, determined by qRT-PCR (the 2-ΔΔCt method). (**D**) A working model of this study, also see *Discussion*. The Kruskal-Wallis test and Dunn’s multiple comparison between all groups were applied to **A**. The Welch’s t-test was applied to **B** and **C**. For all statistical assays, * indicates *p* < 0.05, *** indicates *p* < 0.001, and ‘ns’ indicates not significant. In **A**, all representative images in the same panel share the same scale indicated by a 25 μm scale bar.

Conversely, we wondered if the mere overexpression of one component in this neuropeptide axis would be sufficient to trigger fat mobilization in well-fed wildtype. For this purpose, we generated NLP-10 overexpressed animals and NPR-4 overexpressed animals, but neither was sufficient to elicit fat loss (**Figure 7—figure supplement 1A, B**). This suggests that the neuropeptide signaling pathway is unlikely to serve as an ON/OFF switch for fat mobilization in peripheral tissues, but rather as a signal amplifier that can be modulated by the animal’s nutritional status.

In conclusion, the chronically diet-restricted status produced by either BMP-associated dyspepsia or prolonged food deprivation, could hyperactivate intestinal fat mobilization through a neuropeptide signaling axis involving the NLP-10/NPR-35 and the FLP-18/NPR-4 peptide receptor pairs. Fat loss in diet-restricted animals could be blocked by suppressing any key component — be it a peptide, a receptor, or a lipase. Given sufficient food supply, fat mobilization would require more than the overexpression of a single-factor peptide or receptor.

## DISCUSSION

We demonstrate that diet-restricted *C. elegans* upregulates a neuropeptide signaling axis to stimulate intestinal fat mobilization. This axis involves two peptide-receptor pairs, namely NLP-10-NPR-35 and FLP-18-NPR-4, to enhance triacylglycerol hydrolysis (**Figure 7D**). Rather than a mutant-specific mechanism, this axis generally detects energy deficiency as an internal cue and stimulates fat breakdown as the physiological response.

Starvation-induced neuropeptide upregulation has been characterized across species. In mammals, hypothalamic NPY and AgRP expression increase upon fasting to drive hyperphagia and metabolic adaptation^47,48^. In fasted *Drosophila*, tachykinin rises to suppress lipogenesis^49^. *C. elegans* FLP-18 increases upon food deprivation to modulate food foraging behaviors^50^ and fat metabolism^32^. Our findings have traced the *flp-18*-secreting tissues to the *npr-35-*expressing cells and revealed NLP-10 in ADL neurons as an important upstream signal. The multi-functional ADL neurons have been shown to sense sex pheromones^51,52^ and other stimuli in the external environment, and they can also detect neuronal mitochondrial stress as an internal cue to cause physiological responses in intestine ^53^. In this study, we demonstrate that the ADL neurons are essential for the organism to mobilize fat in response to energy deficiency as an internal stimulus. This echoes and complements previous findings that ADL neurons regulate intestinal fat storage in response to sex pheromones in the external environment ^54^. Considering that the *C. elegans* nervous system is capable of integrating different cues (such as food abundance and pheromone concentration ^55^) to make behavioral decisions, the regulatory effect of ADL neurons and relevant neuropeptides on physiology and decision-making is worthy of further study. In addition to NLP-10, FLP-18, FLP-1 and FLP-4, several peptides have emerged as promising candidates for fat regulation and starvation perception, such as FLP-5, 8, 13, 24 and NLP-6, 9, 13, 35, 50 (**Figure 2— figure supplement 1E**). Elucidating the *in vivo* functions and genetic interactions of these peptides remains challenging yet critical, as such efforts would reveal the operational logic of neuropeptide signaling in modulating metabolic processes—particularly its hierarchical and additive effects on the homeostasis of lipid storage and lipolysis. While many more neuropeptide genes could be transcriptionally induced by nutrient deficit, they are dispensable for regulating fat mobilization (e.g., NLP-15, 21, 51, and FLP-14, 27, as shown in **Figure 7C** and (**Figure 2—figure supplement 1E**), and how these peptides differ from the fat-regulating ones in determining the organism’s metabolic set point remains interesting.

Although transcriptional induction of *nlp-10* has been consistently observed in BMP mutants, *eat-2/4* mutants, and fasted wildtype, the *nlp-10* overexpression *per se* cannot trigger fat mobilization in well-fed animals with intact BMP pathway, nor can NPR-4 overexpression. This suggests that highly redundant mechanisms are programmed in neuroendocrine systems to prevent unnecessary fat mobilization, which permit the organism to store energy and substrates under nutrient-rich conditions. On the flipped side, for the diet-restricted animals to mobilize fat, a series of neuropeptides and receptors need to stay hyperactivated. Would the consistent activation of lipolysis benefit the longevity of BMP mutants? Some studies have shown that the BMP mutants are mildly long-lived ^13,56^. Whether the NLP-10-NPR-35-FLP-18-NPR-4 axis is involved in BMP-associated longevity would require further investigation.

In this study, the importance of including viable and fertile animals with chronic dietary restriction (i.e., BMP mutants) is self-evident for uncovering the detailed action sites and communication patterns of the NLP-10-NPR-35-FLP-18-NPR-4 axis. Now that the dyspepsia phenotype has been identified in the BMP mutants, future investigations can be designed using these chronically dietary-restricted models to address unresolved questions regarding metabolic homeostasis and physiological adaptation. Beyond its traditional bone morphogenetic roles, BMP pathway has been recognized as a multi-faced regulator in lipid metabolism with tissue specificity: In the peripheral tissues of mammals, BMP4 promotes white adipocyte differentiation and has dual roles, either promoting or repressing oxidative metabolism in a cell-context-dependent manner^57,58^; BMP7, in contrast, stimulates brown preadipocyte differentiation to increase energy expenditure and decrease weight gain^59^. In the central nervous system of mammals, BMP signaling also regulates appetite, metabolism and the whole-body energy balance^60^. In *C. elegans*, BMP pathway has been shown to interact with the insulin signaling to regulate lipid accumulation^13^, which is consistent to their chronically diet-restricted state. As for the precise working site of BMP signaling, pharyngeal SMA-3 has been shown to restore intestinal colonization of the pathogenic *Enterobacter* strain^61^, foreshadowing a mechanistic link between BMP signaling and bacteria processing. In wildtype, pharyngeal glands secrete mucin-like proteins to facilitate digestion^62^, and mucins are generally known to mediate a variety of biological processes including oral hydration and non-immune host-defense toward microbes in mammals^63^. Intriguingly, lacking BMP pathway upregulates most mucin-like genes (GO:1905905) ^12^ - whether this is an adaptation to, or a side-effect of the grinding problem merits further investigation.

In *C. elegans* and other models, associations between innate immune responses and BMP signaling are receiving attention ^61,64–66^. Lipidomic analysis of this study reveals that the levels of certain signaling lipids are very different in wildtype and *sma-6(wk7)*. Particularly, eicosanoid, SPH, and ceramide-NP are upregulated, while LPA is downregulated (**Figure 6—figure supplement 1E**), pointing to a coordinated metabolic resetting that dampens reproduction and upregulates immune responses. Specifically, LPA is a bioactive lipid mediator that facilitates germ cell survival and oocyte maturation ^67,68^, eicosanoids serve as CYP-dependent signaling molecules that activate innate immune responses via the p38 MAPK pathway ^69^, and sphingolipids like SPH and ceramide-NP reinforce intestinal epithelial barrier integrity and apical polarity ^70,71^. Thus, the lipidomic profile of *sma-6(wk7)* might reflect a systemic shift from fecundity toward stress defense. Consistently, a category of pathogen-induced genes that are BMP dependent were identified to encode lipid metabolic enzymes ^14^. Whether the shift from fecundity to defense happened to other types of diet-restricted worms would require further investigation.

## SUPPLEMENTARY TABLES

**Supplementary Table1.** Relative gene expression levels by qRT-PCR

**Supplementary Table2**. Intestinal fat levels of animals with BMP deficiency with or without functional neuropeptides

**Supplementary Table 3.** TG-to-DG ratio of wildtype and *sma-6(wk7)* calculated by the sum of area-under-curve values in LC-MS

**Supplementary Table 4.** Reagent and resource used in this study

**Supplementary Table 5.** Oligos used in this study

**Supplementary Table 6.** Raw data for figure plotting

## MATERIALS AND METHODS

### Resources and reagents

*C. elegans* strains were cultured using <u>n</u>ematode <u>g</u>rowth <u>m</u>edia (NGM) agar and *E. coli* OP50 as described ^72,73^, except for the RNAi experiments. For RNAi, bacteria recovered from the Ahringer RNAi library ^74,75^ were fed to *C. elegans* on NGM agar containing 1 mM of Iso<u>p</u>ropyl β-D-1-thio<u>g</u>alactopyranoside (IPTG). All *C. elegans* strains were maintained at 20 °C. **Supplementary table 4** describes the reagents used in this study.

Oligos used in genotyping and plasmid construction were described in **Supplementary table 5**. To construct plasmids, DNA fragments amplified from the genomic DNA or complementary DNA of Bristol N2 *C. elegans* were seamlessly cloned into the pJM23-SL2-GFP backbone. Transgenic strains were generated by standard microinjection methods ^76^ with 30-50 ng/μL of the corresponding plasmids and maintained by picking nematodes with transgenic markers. To verify the expression of *nlp-10p::GFP*, DiI staining ^77^ were applied with the Zeiss Axioplan2 microscopic system at the green channel (525 nm emission wavelength, 470 nm excitation wavelength) and the red channel (610 nm emission wavelength, 560 nm excitation wavelength).

### Starvation and nutrient supplementation

To starve nematodes, synchronized L1s were cultured on NGM agar with *E. coli* OP50 until they reach the L4 stage, rinsed in M9 buffer, and transferred to the NGM agar without bacterial food. After 24 hours of starvation, nematodes were collected as the samples of lipid staining or qRT-PCR experiments. To supplement nematodes with extra nutrients, glucose, fructose or oleic acid was provided to culture synchronized L1s until they reach the adult stage (Day 1). Glucose or fructose were mixed into the NGM agar at 0.5% concentration (w/v). Oleic acid was mixed into the bacterial food at the ratio of 1 : 296 (v/v), and 200 μL of this mixture were provided to each φ = 6 cm NGM agar plate.

### Semi-quantification of intestinal fat levels upon fixation and lipid staining

A Nile Red based lipid staining method, developed from Escorcia et al ^78^ and Stuhr et al ^79^, was applied to determine the relative fat levels: nematodes were briefly rinsed in M9 buffer and immediately fixed in the 1% paraformaldehyde solution for 30 min, followed by a freezing-crack step (frozen at -80 °C for no less than 2 hours) to facilitate stain permeability. Once thawed, the nematodes were rinsed by PBST buffer and 40% isopropanol solution (v/v) before they were soaked into the staining solution (1 ng/μL of Nile Red (MCE® catalog # HY-D0718) in 40% isopropanol) for 30 min. Stained nematodes were rinsed by PBST and mounted onto glass slides with 2% agarose pads, and imaged by the Zeiss Axioplan2 microscopic system at 595 nm emission wavelength and 500 nm excitation wavelength. To quantify fat levels in the intestinal region individual-by-individual, fluorescent intensities were measured by Fiji ImageJ (with the intestinal region being labelled by yellow dotted lines) ^80^. Mean intensity of the control group was defined as ‘1’ for normalization to calculate the relative fat levels.

### Intestinal Colony Forming Units (CFU)

With a method developed from Portal-Celhay et al ^20,81^, the CFU of nematodes’ intestinal bacteria were quantified: Per each genotype, ten adults were picked onto a bacteria-free NGM agar plate containing 100 ug/mL gentamicin (to inhibit *E. coli* growth) to crawl freely for 5 min, and then transferred into a glass vial with M9 buffer containing 25 mM of levamisole (to pause pharyngeal pumping and defecation) and 100 μg/mL gentamicin by glass pipettes, followed by two M9 rinses; 100 μL of PBST were added to the nematode-containing vial, and each vial was manually grinded for 15 sec by a disposable rod; with PBS, the rod was rinsed and a final volume of 500 μL was achieved as the stock solution; the stock solution and its diluted solutions (10^1^ to 10^5^ times) were spread onto LB agar plates to score and calculate CFU per worm upon the 37°C overnight culture.

### Transmission Electron Microscopy (TEM)

Adult nematodes of the same genotype were collected and briefly rinsed in a 1.5mL vial with PBS solution, and 300 μL of ice-cold glutaraldehyde solution (0.67% glutaraldehyde and 0.67% Osmium in 10 mM HEPES) were added to each vial to incubate nematodes for 1 hour on ice, followed by 5 rinses of ice-cold 10 mM HEPES (15 min per each rinse). The nematodes were then fixed in ice-cold OsO_4_ solution (2% osmium in 10 mM HEPES) and incubated for 3 hours at 4°C. Another 5 rinses (by water) were applied before the nematodes were embedded in 2% low melt agarose and stained in 1 % uranyl acetate overnight. Upon a brief rinse in water, samples were dehydrated in a cold graded ethanol series (20 %, 50 %, 70 %, 90 %, 100 %, 100 %) for 13 min per each, followed by two acetone rinses at room temperature, and infiltrated in SPI-Pon 812 using anhydrous acetone-to-812 at different volume ratios: 3:1, 1:1, 1:3, and then 100 % of 812 overnight. Before baking at 60 °C for 48 hours, one more rinse with fresh 812 were applied. The embedded nematodes were then sectioned into 75 nm slices using the Leica UC7 ultramicrotome, stained by lead citrate for 5 min, and imaged by the Hitachi HT7800 transmission electron microscope.

### Differentially expressed genes (DEGs) and Gene Ontology (GO) analysis

RNA-seq data generated in Yu et al ^12^ were used for the identification of DEGs and GO analysis by Metascape at https://metascape.org/ ^82^.

### Lipidomic profiling and data analysis

Synchronized wildtype (Bristol N2) and *sma-6(wk7)* young adults (∼10,000 nematodes per genotype per replicant) were collected by M9 buffer, followed by removal of supernatant and immediate frozen via dry ice. Lipid extraction, LC-MS detection, lipid identification and quantification, and data analysis were purchased from MetwareBio Inc. To calculate the TG-to-DG ratio, corresponding area-under-curve values were summed and standardized as the dividends and divisors.

### Statistical analysis and scientific plotting

Unless otherwise indicated, statistical significance was assessed using a two-tailed Mann-Whitney test to compare two genotypes, or Kruskal-Wallis test to compare more than two genotypes. Scientific plotting was achieved by GraphPad Prism 10.1 and Adobe Illustrator 2023.

## ACKNOWLEDGEMENTS

For reagents, some strains were provided by the CGC, which is funded by NIH Office of Research Infrastructure Programs (P40 OD010440), and some strains were provided by the NBRP, which is funded by the Japanese government. We thank Prof. Meng C Wang at HHMI Janelia, for kindly providing some reagents and offering advice. We thank Prof. Douglas S. Portman at University of Rochester for kindly offering suggestions to this study. We thank Prof. Geng Wang at Xiamen university for suggestions. We thank all graduate students, undergraduate students and alumni in Yu Lab at Xiamen University for technical supports. This work was supported by the National Natural Science Foundation of China (32071146) and the Fujian Provincial Natural Science Foundation of China (2024J01025) to Y.Y.; and by the Fundamental Research Funds for the Central Universities (20720230066) and the Fujian Provincial Natural Science Foundation of China (2023J05007) to J.L.

## AUTHOR CONTRIBUTIONS

Conceptualization, Y. Yu, J. Liu and J. Luo; Investigation, J. Liu, L. Lin, L. Yao, J. Luo and Y. Yu; Writing (original draft), J. Luo, J. Liu, L. Lin and Y. Yu; Funding acquisition, Y. Yu and J. Luo.

